# Identification of single pair of interneurons for bitter taste processing in the *Drosophila* brain

**DOI:** 10.1101/170464

**Authors:** Ali Asgar Bohra, Benjamin R Kallman, Heinrich Reichert, K. VijayRaghavan

## Abstract

*Drosophila* has become an excellent model system for investigating the organization and function of the gustatory system due to the relatively simple neuroanatomical organization of its brain and the availability of powerful genetic and transgenic technology. Thus, at the molecular and cellular level, a great deal of insight into the peripheral detection and coding of gustatory information has already been attained. In contrast, much less is known about the central neural circuits that process this information and induce behaviorally appropriate motor output. Here we combine functional behavioral tests with targeted transgene expression through specific driver lines to identify a single bilaterally homologous pair of bitter sensitive interneurons that are located in the subesophageal zone of the brain. Anatomical and functional data indicate that these interneurons receive specific synaptic input from bitter sensitive gustatory receptor neurons. Targeted transgenic activation and inactivation experiments show that these bitter sensitive interneurons can largely suppress the proboscis extension reflex to appetitive stimuli such as sugar and water. These functional experiments together with calcium-imaging studies indicate that these first order local interneurons play an important role in the inhibition of the proboscis extension reflex that occurs in response to bitter tastants. Taken together, our studies present a cellular identification and functional characterization of a key gustatory interneuron in the bitter sensitive gustatory circuitry of the adult fly.

## INTRODUCTION

The gustatory and olfactory systems of *Drosophila* represent powerful models for analyzing the neuronal organization of chemosensory systems (Vosshall and Stocker, 2007). In the adult olfactory system, a great deal is now known about the structure and function of the circuitry that detects and processes olfactory information (Masse *et al.*, 2009; Hong and Luo, 2014; Joseph and Carlson, 2015); There are approximately 50 different classes of olfactory sensory neurons, each with a specific olfactory receptor type. Each olfactory sensory neuron type projects its axon to a single glomerulus in the antennal lobe of the brain where synaptic interactions with local interneurons and projection neurons take place. Projection neurons convey processed sensory information from the glomeruli to higher order brain centers in the mushroom bodies and lateral horn which process olfactory information further for behavioral functions such as learning and memory or appetitive and aversive response control.

In the adult gustatory system, considerable insight into the molecular and cellular mechanisms of taste perception has also been attained. Functionally distinct classes of gustatory receptor neurons (GRNs) have been identified including GRNs for bitter tastants such as those labeled by the GR66a receptor and GRNs for sweet tastants such as those labeled by the Gr5a and Gr64f receptors (Thorne *et al.*, 2004; Wang *et al.*, 2004; Marella *et al.*, 2006; Dahanukar *et al.*, 2007; Jiao *et al.*, 2008; Weiss *et al.*, 2011); GRNs for salt, water and pheromone detection have also been identified (Liman, Zhang and Montell, 2014; Freeman and Dahanukar, 2015; Joseph and Carlson, 2015). Moreover, and the gustatory receptor molecules expressed in these different GRN classes have also been characterized (see Freeman and Dahanukar, 2015). Tastant-driven GRN activation results in modality-specific behavioral responses. Thus, activation of sweet GRNs stimulates feeding behavior such as the proboscis extension reflex (PER) and ingestion while activation of bitter GRNs promotes aversive behavior such as PER inhibition and avoidance of noxious compound mixtures (Dethier, 1976; König *et al.*, 2014; French *et al.*, 2015).

The axons of the GRNs, which are located on the proboscis, legs and wing margins, project into discrete regions of the subesophageal zone (SEZ) in the central brain, which is the initial processing center for gustatory information (Stocker, 1994; Vosshall and Stocker, 2007; Ito *et al.*, 2014). The SEZ also comprises the motor neurons, which control the proboscis extension reflex (PER) that occurs in response to appetitive gustatory sensory input, and motor neurons that control ingestion (Rajashekhar and Singh, 1994; Gordon and Scott, 2009; Hampel *et al.*, 2011; Manzo *et al.*, 2012; Schwarz *et al.*, 2017). Moreover, the SEZ also contains a pair of local interneurons that have command function in the feeding motor program (Flood *et al.*, 2013). Thus, local circuits might exist within the SEZ that mediate the transformation of the appropriate sensory input from GRNs into the proboscis motor neuron output required for feeding. In support of this notion, recent large-scale calcium imaging studies indicate that approximately 70 neurons located in the SEZ respond to either sweet or bitter gustatory input and that the majority of these cells are not motor neurons but rather modality-specific interneurons (Harris *et al.*, 2015). Interestingly, these studies also suggest that the gustatory input to motor neurons is similarly modality-specific (sweet and bitter tastants activate different motor neurons) implying that neural circuitry for sweet and bitter gustatory information processing in the SEZ is largely segregated from sensory input to motor output.

Given the large number of interneurons in the SEZ that appear to respond to gustatory input, it is remarkable that only very few of these have been identified at the cellular level. Three gustatory interneuron types that respond to sweet taste input have been identified in the adult SEZ (Kain and Dahanukar, 2015; Yapici *et al.*, 2016; Kim, Kirkhart and Scott, 2017). All are thought to receive direct synaptic input from sweet sensitive GRNs implying that they are first order gustatory interneurons. In contrast, information about first order interneurons that receive and process gustatory information about other tastant categories such as bitter, salt and water is largely lacking. This lack of information has been a major stumbling block for unraveling gustatory circuits in the SEZ and understanding the interneuronal processing of different taste modalities in the gustatory system of the adult fly.

Here we use functional behavioral screening with transgenic driver lines to identify and characterize bitter sensitive gustatory interneurons in the adult SEZ. By screening Gal4 lines for their ability to inhibit sugar-induced appetitive PER by targeted transgenic activation via TrpA1, we identify the VGN6341 line as a marker for interneurons implicated in aversive gustatory responses. Further experiments that limit the targeted transgenic activation to specific CNS regions show that the corresponding VGN6341 interneurons are located in the SEZ. GFP reconstitution across synaptic partners (GRASP) experiments indicate that these interneurons receive direct synaptic input from Gr66a labeled bitter-sensitive GRNs. Combination of VGN6341 cis-flipout with functional behavior assay identifies a single bilaterally symmetric pair of SEZ interneurons as the first order gustatory interneurons responsible for the inhibition of the appetitive PER. Further activity assays using calcium imaging provide functional confirmation for the existence of the VGN6341 interneuron pair in the SEZ that is activated by natural or transgenic stimulation of bitter GRNs.

## RESULTS

### Identification of candidate neurons involved in aversive taste behaviors

To identify candidate neurons that might be involved in gustatory circuitry, we performed a functional behavioral screen, in which Gal4 driver lines were used together with a UAS::TrpA1 reporter to activate target neurons, and the resulting effects on sucrose-induced appetitive PER was examined (Figure 1A). We reasoned that Gal4 driver lines that were able to inhibit the tastant-induced PER through targeted TrpA1 expression (Gal4>TrpA1) might be potential markers for interneurons involved in aversive bitter responses. (TrpA1 is a heat-activable cation channel that can be used to depolarize the neurons in which it is expressed; Hamada et al., 2008). This screen resulted in the identification of the VGN6341 Gal4 line as a driver that targets putative candidate interneurons of the gustatory bitter processing circuitry.

**Figure 1.**
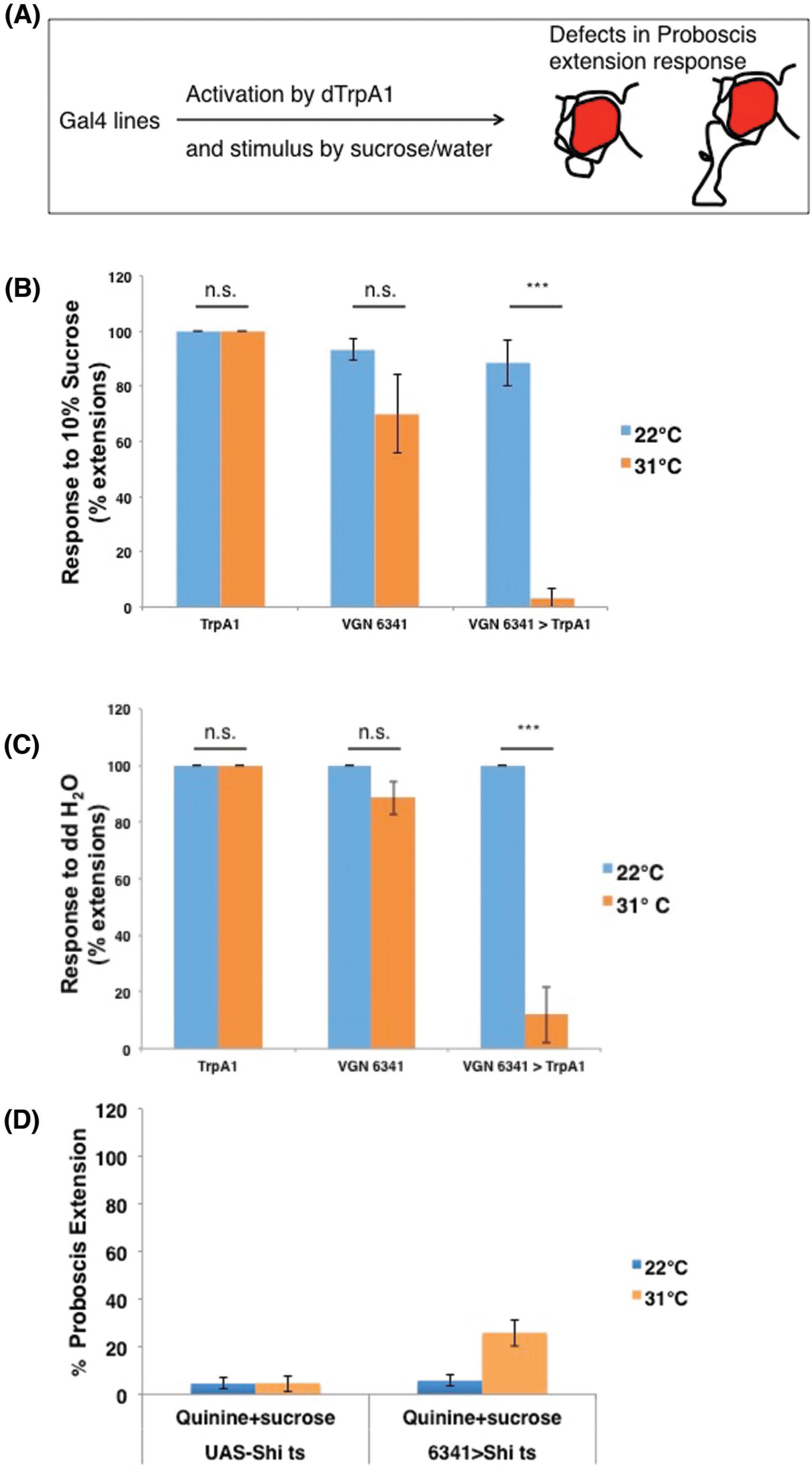
VGN6341 labels neurons involved in inhibition of appetitive behavior. (A) Schematic for behavioral screening by activating neurons in selected GAL4 lines. (B) Quantification of PER phenotypes upon activation of VGN 6341 GAL4 neurons and stimulation with 10% sucrose. (C) Quantification of PER phenotypes upon activation of VGN 6341 GAL4 neurons and stimulation with water. (D) Quantification of PER phenotypes upon silencing of VGN 6341 GAL4 neurons and stimulation with 10% sucrose+lmM quinine mixture.*p < 0.05, **p < 0.01, ***p < 0.001, Student’s *t* test was used for all statistical comparisons. For each bar n=4 trials of 15 flies each.

In non-satiated wild type flies, tarsal stimulation with sucrose reliably elicits appetitive PER; sugar stimulation results in one or more PER in virtually to 100% of the trials (Dethier, 1976; Wang *et al.*, 2004; Masek and Scott, 2010). Like wild type flies, control flies carrying either TrpA1 or VGN6341 reliably extended their proboscis in response to tarsal sucrose stimulation, and they did so at both 22°C and 31°C. Moreover flies carrying both VGN6341 and TrpA1 (VGN6341>TrpA1) also reliably extended their proboscis in response to tarsal sucrose stimulation at 22°C (TrpA1 inactive). However, these flies rarely extended their proboscis in response to tarsal sucrose stimulation at 31°C (TrpA1 active); PER occurred in response to sugar stimuli in less than 4% of the trials (Figure 1B) (supplementary movie 1).

Comparable results were obtained in experiments in which the PER was elicited in thirsty flies by tarsal stimulation with water. Control flies carrying either TrpA1 or VGN6341 reliably extended their proboscis in response to tarsal water presentation at both 22°C and 31°C. Flies carrying both TrpA1 and VGN6341 (VGN6341>TrpA1) also reliably extended their proboscis in response to tarsal stimulation with water at 22°C. However, they rarely extended their proboscis in response to tarsal stimulation with water at 31°C (Fig. 1C). These findings indicate that transgenic activation of VNG6341 neurons can result in inhibition of the PER to an appetitive gustatory stimulus such as sucrose or water.

In view of these findings, we wondered if inactivation of VGN6341 neurons might have an opposing effect on the PER. To investigate this, we took advantage of the fact that the PER to an appetitive stimulus such sucrose can be largely eliminated if sucrose is presented together with an aversive stimulus such as quinine e.g. (König *et al.*, 2014; French *et al.*, 2015). For targeted inhibition of the VGN6341 neurons we used a temperature sensitive UAS::Shi^ts^ reporter. (At restricted temperatures, Shi^ts^ silences neurons by preventing synaptic vesicle reuptake; (Kitamoto, 2001) As expected, at 22°C (permissive temperature) and 31°C (restrictive temperature), control flies carrying the Shi^ts^ reporter (UAS-Shi^ts^) rarely extended their proboscis in response to a sucrose/quinine combination. Flies carrying both Shi^ts^ and VGN6341 (VGN6341>Shi^ts^) also rarely extended their proboscis in response to the sucrose/quinine combination at 22°C; PER occurred in only 5% of the trials. However, these flies showed a significant increase in PER to the aversive sucrose/quinine stimulus mixture at 31°C; PER occurred in 25% of the trials (Fig. 1D). As the PER increase is less than sugar controls, the neurons in VGN6341 are unlikely to be the only neurons that convey bitter taste signals.

Taken together, these results indicate that activation of VGN6341 neurons can inhibit the PER response to an appetitive gustatory stimulus such as sucrose or water and that inactivation of VGN6341 neurons can enhance the PER response to an aversive gustatory stimulus mix such as a sucrose/quinine combination. Thus, activating VGN6341 neurons has a similar behavioral effect as activating bitter sensitive gustatory circuitry via bitter GRN stimulation. Moreover, inactivating VGN6341 neurons has a similar behavioral effect as inactivating bitter sensitive gustatory circuitry via bitter GRN inhibition. To determine if the VGN6341 neurons might indeed comprise neural elements of the bitter sensitive gustatory circuitry, we first characterized these neurons further anatomically using Gal4/UAS labeling experiments.

### VGN6341 Gal4 labels neurons in the central brain and ventral nerve cord

The neuroanatomical features of the VGN6341 neurons can be revealed in targeted label experiments by using VGN6341 Gal4 to drive UAS mCD8::GFP. These experiments show that both the central brain and the ventral nerve cord contain subsets of labeled neurons.

In the central brain, the majority of the labeled neurons are located in the subesophageal zone (SEZ); only sparse cell labeling is seen more anterior brain regions (Fig 2A). The labeled neurons in the SEZ have cell bodies located in two small clusters on each side of the midline, one medial and one lateral (Fig. 2A). In order to map the putative presynaptic terminals of these neurons, we expressed the presynaptic marker synaptotagmin-hemagglutinin (HA) in these neurons (Robinson *et al.*, 2002). Immunolocalization with α-HA showed extensive labeling in the medial SEZ (Fig 2B). The synaptotagmin labeling thus implies that majority of these labeled neurons are interneurons that do not project axons out of the CNS. In addition to these interneurons, 3 labeled motor neurons are also present in the SEZ; these motor neurons innervate 3 different muscles of the proboscis (Fig 2C).

**Figure 2.**
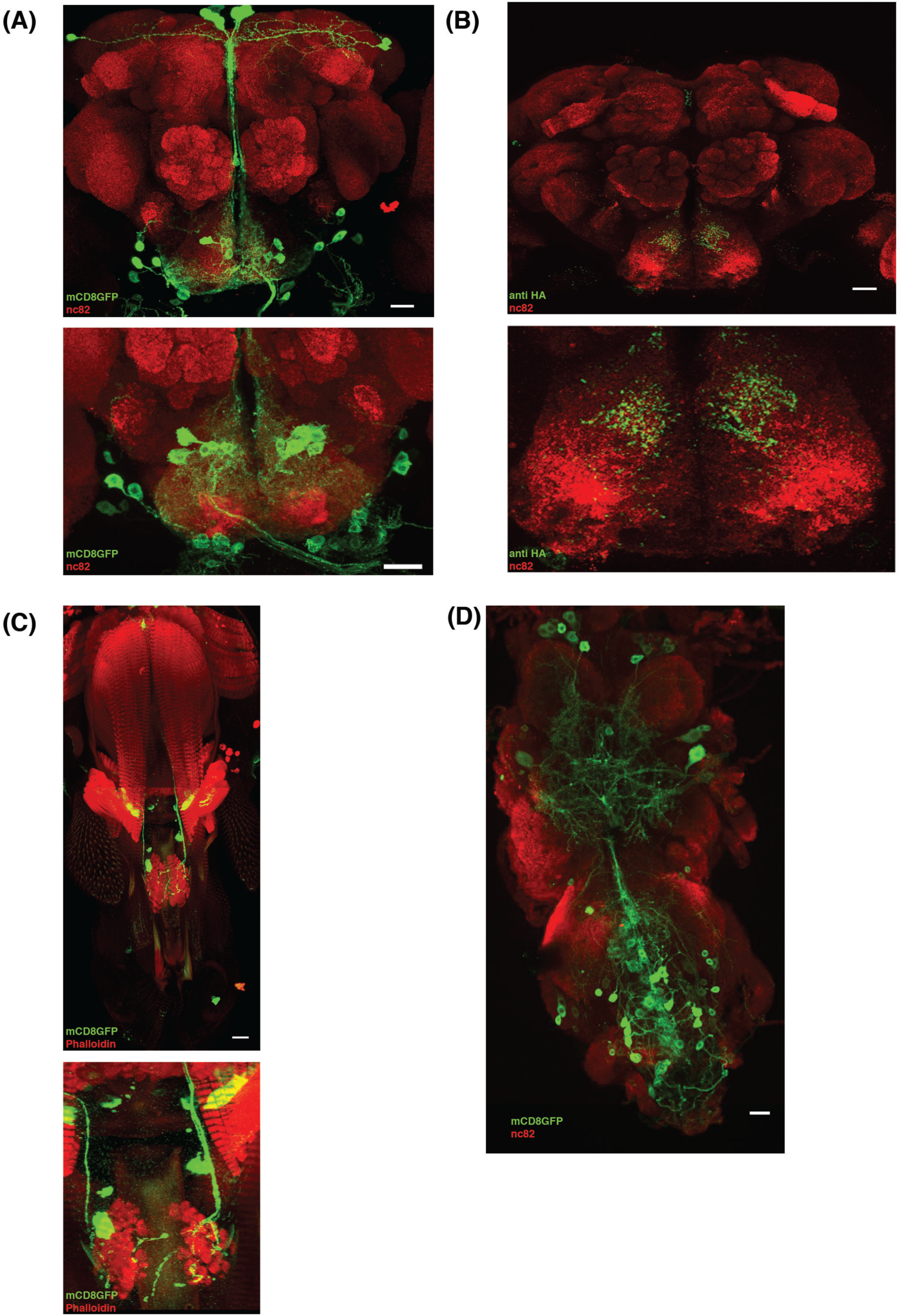
Expression pattern of VGN 6341 GAL4 in the central nervous system. (A) VGN 6341-GAL4-driven expression of UAS-mCD8::GFP in the whole brain (upper) and zoomed in SEZ (lower) shows expression predominantly in approximately 20 cells in SEZ region. (B) Visualization of pre synaptic terminal using flies of VGN 6341 Gal4/+; UAS syt-HA/+ shows the projection of VGN 6341 neurons in the whole brain (upper) and zoomed in SEZ. (C) Motor neurons innervating proboscis muscles 2,5 and 8 are labeled by VGN 6341, shown in upper panel and zoomed in lower panel. (D) VGN 6341-GAL4 driven expression of UAS-mCD8::GFP in ventral nerve cord show expression in thoracic and abdominal ganglion. Scale Bar, 20 μm

In the ventral ganglia, labeled neurons are seen in both the thoracic and the abdominal neuromeres (Fig. 2D). These labeled cells include numerous leg motor neurons (Syed *et al.*, 2016). In contrast, neither GRNs in the proboscis (Supplementary Fig.1) nor GRNs in the leg tarsi are labeled by VGN6341 (data not shown).

Given the regionally diverse distribution of neurons labeled by VGN6341 Gal4, we next determined which of these neuronal subsets was responsible for inhibiting the PER to appetitive gustatory stimuli.

### VGN6341 interneurons in the SEZ are responsible for the inhibition of appetitive PER

To investigate if the VGN6341 neurons located in the ventral ganglia are required for the inhibition of appetitive PER, we limited the transgenic activation via TrpA1 to the neurons in the brain. For this, VGN6341 Gal4 was used to drive UAS-TrpA1 in a Tsh Gal80 background. Tsh Gal80 inhibits the Gal4/UAS system in neurons of VNC, but has no effect on the Gal4/UAS system in the central brain (Clyne and Miesenböck, 2008) (Fig. 3A, B). In response to tarsal sucrose stimulation, control flies carrying Tsh Gal80 and TrpA1 but lacking VGN6341 reliably extended their proboscis at both 22°C and 31°C. In contrast, flies carrying VGN6341 and TrpA1 (VGN6341>TrpA1) in the presence of Tsh Gal80 reliably extended their proboscis in response to tarsal sucrose stimulation at 22°C (TrpA1 inactive), but rarely extended their proboscis in response to tarsal sucrose stimulation at 31°C (TrpA1 active) (Fig 3C). This finding indicates that VGN6341 neurons in the ventral ganglia are not required for the inhibition of the appetitive PER.

**Figure 3.**
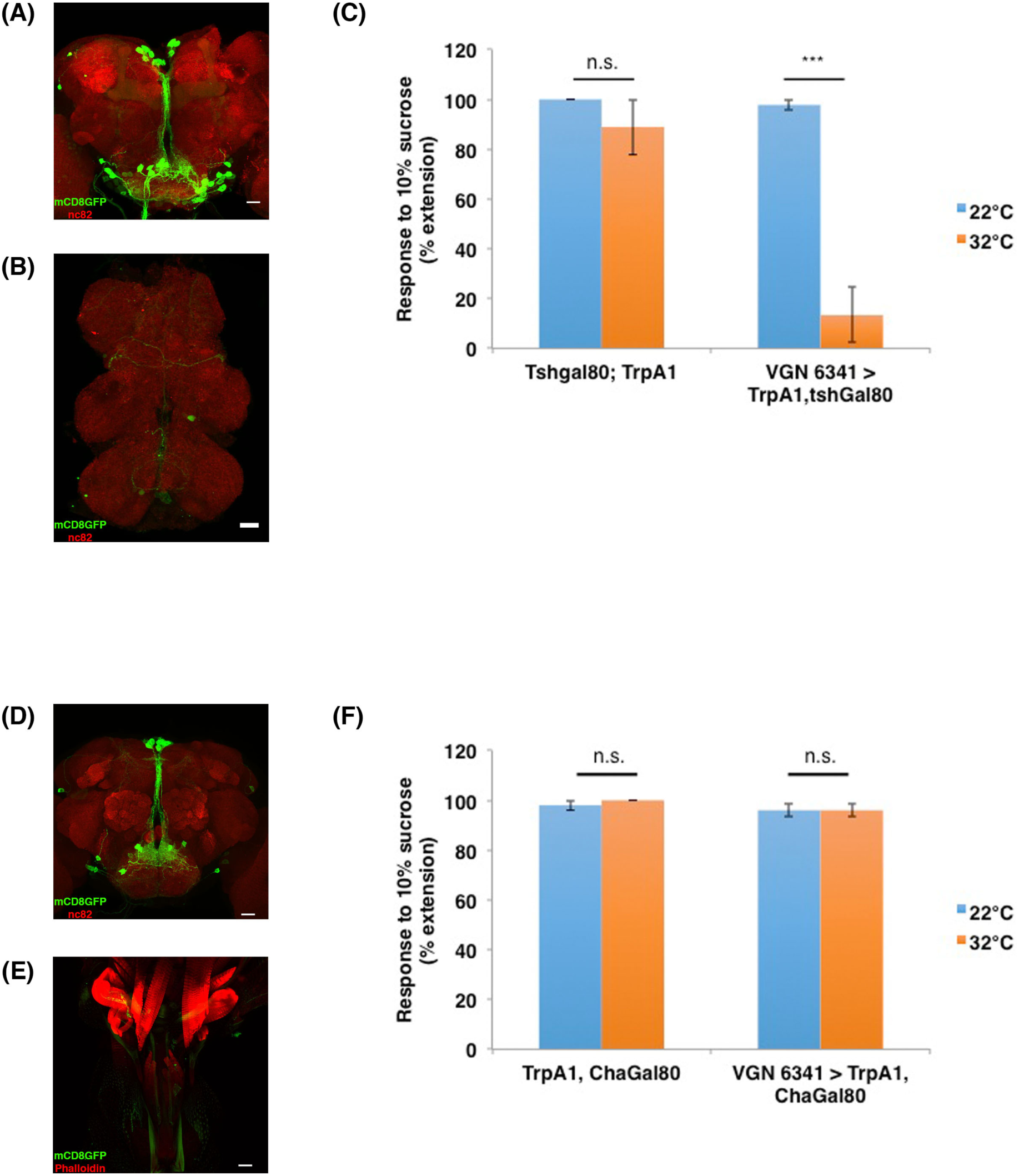
Interneurons labeled by VGN 6341 in central brain are responsible for inhibition of appetitive PER. (A) Expression of mCD8::GFP driven by VGN 6341 GAL4; tsh-GAL80 in central brain remains unchanged. (B) Tsh-GAL80 removes the labeling of neurons by VGN-6341-GAL4 in ventral nerve cord. (C) Quantification of PER phenotypes upon activation of VGN 6341 GAL4 neurons in background of tsh-Gal80 and stimulation with 10% sucrose. Left side control without VGN 6341 Gal4 driver, right side experimental with VGN 6341 Gal4 driver. (D) Expression of mCD8::GFP driven by VGN 6341 GAL4;Cha-GAL80 in central brain marks motor neurons in SEZ and few interneurons in higher brain. (E) Cha-GAL80 does not remove labeling of motor neurons via VGN 6341 GAL4 driver and hence motor neuron innervation to proboscis muscles remains unchanged. (F) Quantification of PER phenotype upon activation of VGN 6341 GAL4 neurons in presence of Cha-Gal80 and stimulation with 10% sucrose Left side control without VGN 6341 Gal4 driver, right side experimental with VGN 6341 Gal4 driver. *p < 0.05, **p < 0.01, ***p < 0.001. For each bar n= 3 trials of 10 flies each.

The population of VGN6341 neurons in the brain consists of both interneurons and motor neurons (see above). To rule out that VGN6341 motor neurons in the brain (or elsewhere) are responsible for the induced inhibition of appetitive PER, VGN6341 Gal4 was used to drive UAS-TrpA1 in the presence of Cha Gal80, which inhibits the Gal4/UAS system in cholinergic interneurons but not in the (glutamatergic) motor neurons (Fig. 3D, E). In response to tarsal sucrose stimulation, control flies carrying Cha Gal80 and TrpA1 but not VGN6341 reliably extended their proboscis at both 22°C and 31°C. Flies carrying VGN6341 and TrpA1 (VGN6341>TrpA1) in the presence of Cha Gal80 also reliably extended their proboscis in response to tarsal sucrose stimulation at both 22°C and 31°C (Fig 3F). This finding indicates that the motor neurons targeted by VGN 6341 are not responsible for the inhibition of the appetitive PER.

Taken together, these experiments indicate that the PER behavioral inhibition elicited by VGN6341 Gal4-driven activation is due to VGN6341 interneurons present in the SEZ of the brain. However, these findings do not indicate if these relevant interneurons are elements of the bitter sensitive gustatory circuitry.

### Bitter sensory neurons show GRASP with VGN6341 neurons in the SEZ

To investigate if some of the VGN6341 interneurons in the SEZ might receive synaptic input from bitter GRNs we carried out GRASP split GFP labeling. The GRASP (GFP reconstitution across synaptic partners) technique reveals if processes of two neuronal populations labeled with different drivers lie in close proximity to each other as would be expected for synaptic partners (Feinberg *et al.*, 2008; Gordon and Scott, 2009). For this, one half of the split GFP reporter (lexAop-spGFP11) was targeted to bitter sensitive GRNs using Gr66a-LexA as a driver, and the other half of the split GFP reporter (UAS-spGFP1-10) was targeted to the VGN6341 neurons using VGN6341-Gal4 as a driver. GRASP visualization was using an antibody that does not efficiently recognize either of the split GFP fragments alone.

Visualization of the sensory neurons labeled by Gr66a-LexA driving a full length GFP reporter revealed the axons of the bitter GRNs as well as the morphology of their terminal arbors in the appropriate zones of the SEZ as shown previously (Figure 4A; Wang *et al.*, 2004). Visualization of the neurons labeled by VGN6341-Gal4 driving a full-length GFP reporter revealed the morphology of the labeled interneurons and motor neurons in the SEZ (Fig. 4B). No labeling was seen in the SEZ in GRASP control experiments in which the VGN6341-Gal4 driver was combined with the UAS-spGFP1-10 reporter (Fig. 4C). In contrast, when GRASP between these two neuronal populations was visualized by immunolabeling of reconstituted GFP, a clear GFP signal was observed in the region of the SEZ where which Gr66a GRN axons terminate (Fig 4D).

**Figure 4.**
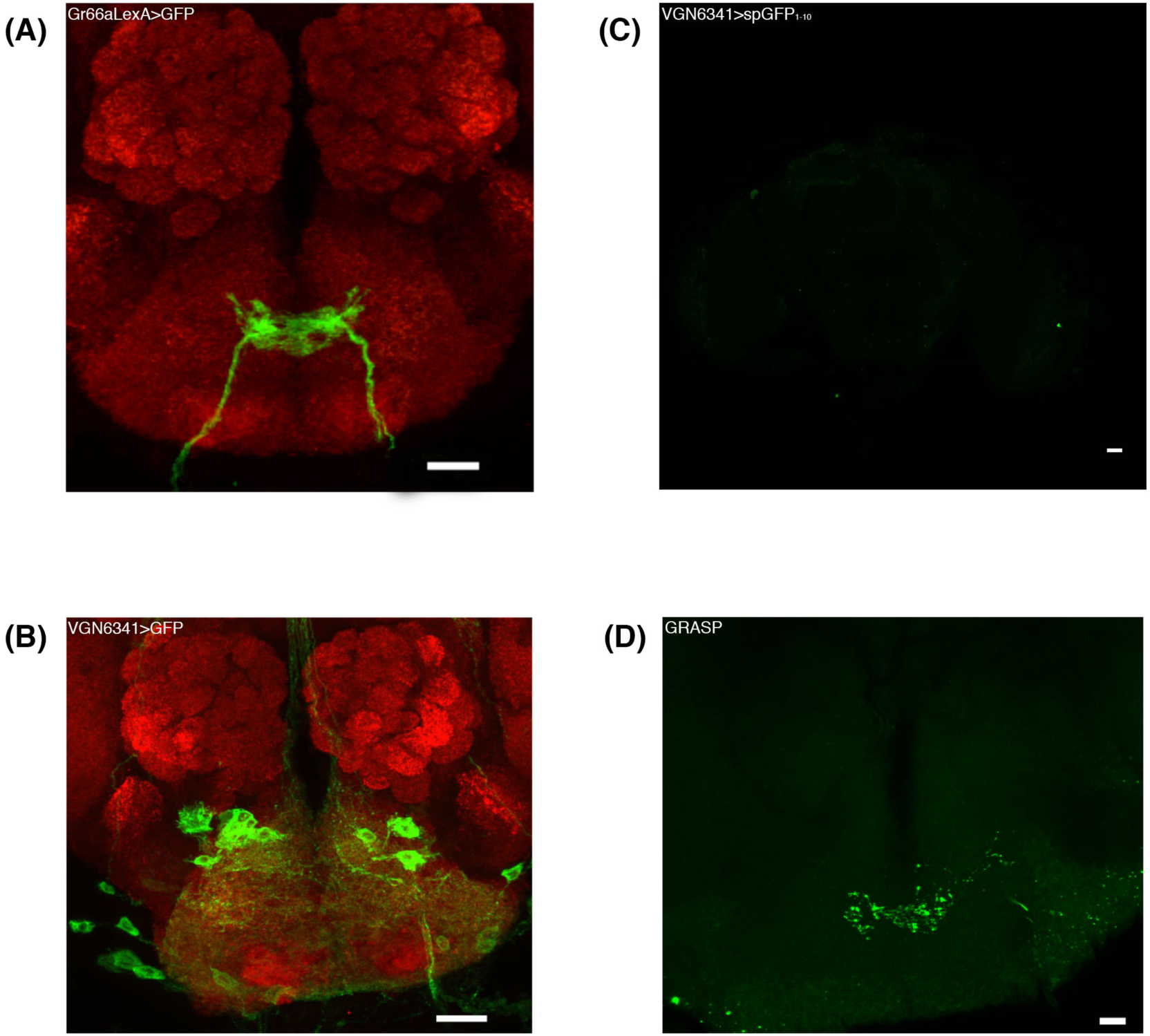
VGN 6341 neuron synapse with bitter sensory neurons. (A) Axonal termini of the bitter sensory neurons in SEZ marked by flies of genotype Gr66a LexA; LexAop mCD8GFP. (B) Expression pattern of VGN 6341 labeled neuron in SEZ showing dendritic arbors of neurons in proximity of axonal termini of bitter sensory neurons. (C) Image of the SEZ with α GFP of control flies with genotype VGN 6341>UAS-CD4::SpGFP1-10. (D) Representative image of the SEZ showing GRASP signal in flies with genotype Gr66a LexA/LexA-op-CD4::SpGFP11;VGN 6341/UAS-CD4::SpGFP1-10 (n=10). Scale bars, 20 μm

This finding indicates that the terminals of Gr66a bitter sensitive GRNs are in close proximity to processes of the VGN6341 neurons in the SEZ. This in turn suggests that at least one of the VGN6341 neurons receives synaptic input from bitter GRNs and, thus, corresponds to a first order bitter sensitive interneuron that, hence would be part of the bitter sensitive gustatory circuitry.

### Identification of a VGN6341 interneuron sufficient for the inhibition of appetitive PER

In order to identify the postulated bitter sensitive interneuron(s) in more detail, we combined VGN6341 cis-flipout with a functional behavioral assay in a genetic mosaic analysis designed to identify interneurons responsible for the inhibition of the appetitive PER at the single cell level. For this, heat shock was used to flip out GAL80 (tub-FRT-GAL80-FRT) and induce VG6341-dependent expression of UAS-dTrpA1 and UASmCD8::GFP in the SEZ. We recovered a total of 66 flies with small sets of labeled VG6341 neurons in the SEZ. Among these, 21% manifested an inhibition of appetitive PER at 31°C (TrpA1 active) and 79% did not.

In the flies that did show an inhibition of appetitive PER, the VG6341-labeled neurons always comprised a pair of bilaterally symmetric SEZ interneurons, either in isolation or together with other neurons. Representative images of these interneurons are shown in Fig. 5A. Examples of images of neurons that did not show inhibition of appetitive PER are shown in Fig. 5B. These findings suggest that the activation of a single identified SEZ interneuron pair among the population of VGN6341 neurons is sufficient for the inhibition of the appetitive PER.

**Figure 5.**
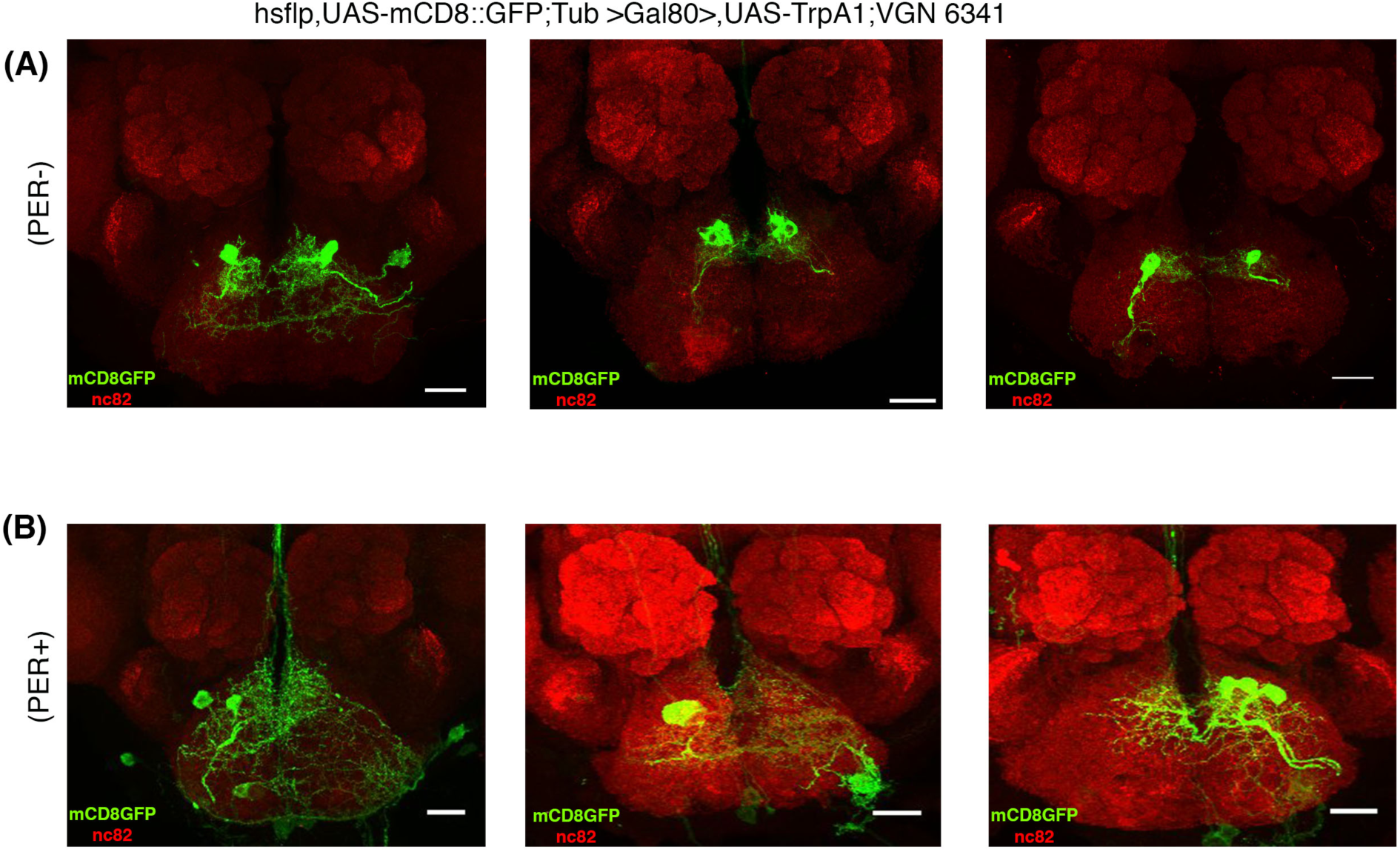
Clonal analysis reveals a pair of interneuron in SEZ region that Mediate inhibition of PER. (A) Representative images of GFP+ clones of pair of interneurons in the SEZ mediating inhibition of proboscis extension response. Clones are derived from hsflp, UAS-mCD8::GFP;Tub >Gal80>,UAS-TrpA1;VGN 6341 flies. n=14 (B) Flies showing normal proboscis extension response don’t contain the interneurons mediating inhibition of PER. Scale bar, 20 μm; n=52.

In anatomical terms, this interneuron has its cell body located in the medial cluster of VGN6341 labeled cells and forms highly branched dendrite-like processes in the region of the SEZ neuropil in which Gr66a sensory axons terminate. Moreover it projects a single short unbranched axon-like process towards centromedial SEZ neuropil where it forms terminal arbors. Since none of the interneuron’s processes project out of the SEZ, the labeled cell has the morphological features of a SEZ local interneuron.

Based on its anatomical characteristics and considering its functional effects on appetitive PER, we postulate that this VGN6341 interneuron corresponds to a first-order bitter sensitive gustatory local interneuron (bGLNs). If this is the case, activation of bitter sensitive GRNs through bitter tastants or through transgenic targeted activation of bitter sensitive GRNs should lead to the activation of a single, medial cluster VGN6341 interneuron in each half of the SEZ. To investigate this, we carried out calcium-imaging experiments designed to monitor the activity of neurons in the SEZ at single cell resolution.

### Calcium imaging reveals functional connections of bitter GRNs and the VGN6341 local gustatory interneuron

To monitor the response of VGN labeled interneurons of the SEZ, we expressed GCaMP6s, a genetically encoded calcium indicator with high sensitivity under the control of the VGN6341-Gal4 driver (Chen *et al.*, 2013; Harris *et al.*, 2015). To elicit gustatory input, proboscis stimulation with a potent bitter tastant combination (denatonium/caffeine/polyethylene glycol; see methods) or transgenic activation of Gr66a-labeled afferents via heat-activated TrpA1 was carried out in the living fly. The resulting changes in fluorescence in the SEZ were acquired with spinning disk confocal microscopy (see methods; Harris *et al.*, 2015).

Gustatory stimulation of the proboscis with the bitter tastants and simultaneous calcium imaging revealed changes in fluorescence in the cell body of a single (VGN6341-specific) interneuron in each half of the SEZ (Fig. 6A) (Supplementary movie 2). The affected cell body was consistently located in the medial cluster of VGN6341 neurons. Transgenic activation of the bitter sensitive Gr66a GRNs using TrpA1 and simultaneous calcium imaging also resulted in changes in fluorescence in the cell body of a single interneuron in each half of the SEZ (Fig. 6B) (Supplementary movie 3). Moreover, the location of the affected cell bodies was the same as in the experiments involving proboscis stimulation. In contrast, control proboscis stimulation with sucrose did not elicit a significant fluorescence change in the SEZ (Fig. 6C).

**Figure 6.**
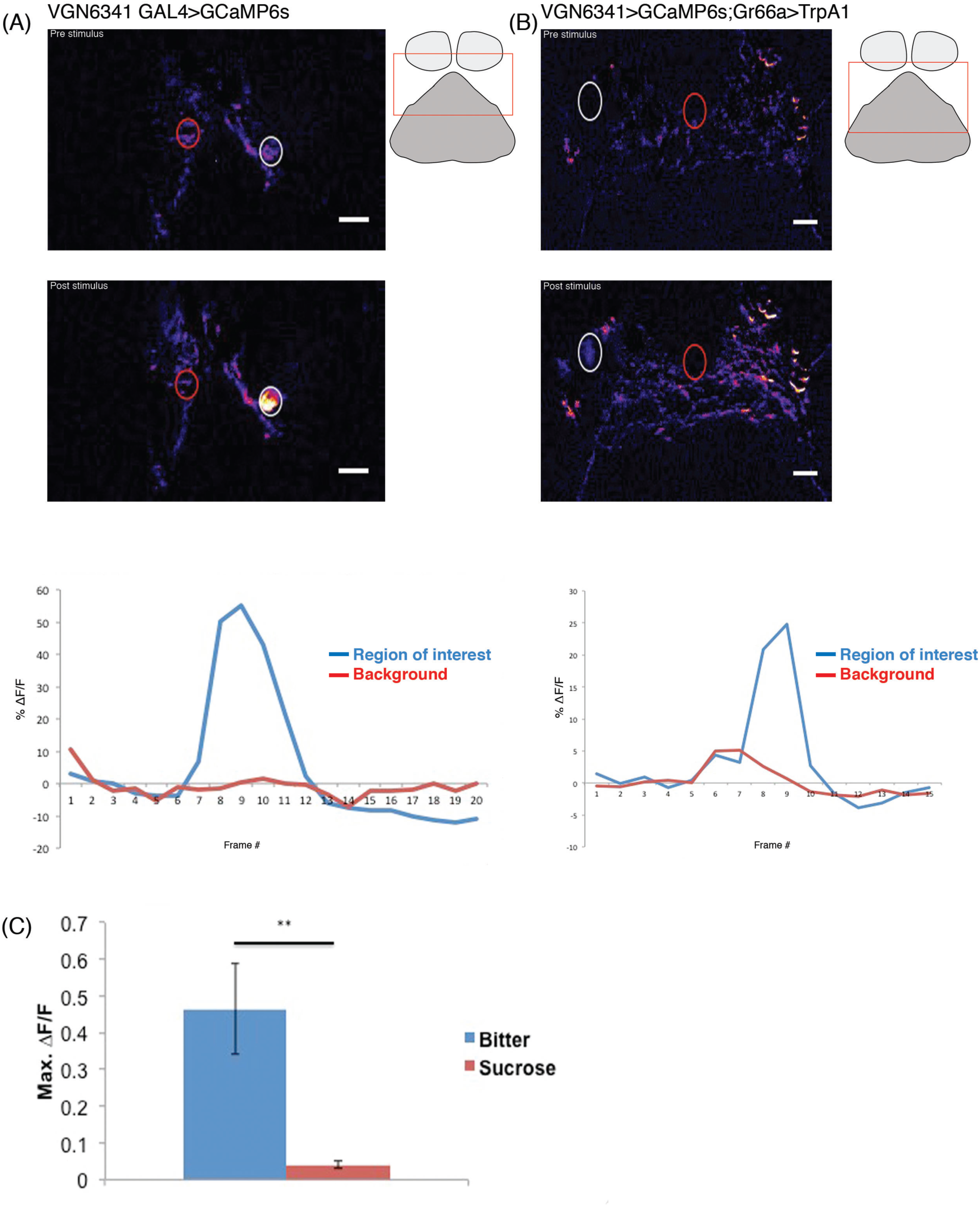
VGN 6341 bGLNs are activated by bitter tastants. (A) Color coded images and mean fluorescence change in VGN 6341 bGLNs. The top image is pre-stimulus change in GCaMP fluorescence; middle image is post-stimulus showing change in fluorescence (ΔF/F) to bitter stimulant. Representative traces showing fluorescence changes of VGN 6341 bGLNs in response to stimulation with bitter (bottom). Schematic showing approximate region of calcium imaging (rectangular box). (B) Color coded images and mean fluorescence changes of GCaMP in VGN 6341 bGLNs in flies of the genotypes VGN 6341 > GCaMP6s; Gr66a > dTrpA1. The top image is before heat activation of Gr66a^+^ neurons by IR laser; middle image is following heat activation by IR laser showing change in fluorescence (ΔF/F). Representative traces showing fluorescence changes of VGN 6341 bGLNs in response to heat activation of bitter sensory neurons (bottom). Fluorescence intensity measurements were taken from areas within dashed circles: region of interest (white) and background (red). Schematic showing approximate region of calcium imaging (rectangular box). (C) Max fluorescence changes of GCaMP in VGN 6341 bGLNs upon stimulation of the proboscis with bitter and sugar tastants. (n=6-8 flies for each tastants). **p < 0.01.

Taken together, these experiments demonstrate the existence of functional connections between bitter sensitive GRNs and a single interneuron (pair) in the medial cluster of VGN6341 neurons in the SEZ. These functional data from calcium imaging studies support the findings of anatomical and behavioral studies that identify the interneuron in the medial cluster of VGN6341 neurons in the SEZ as a first-order bitter sensitive gustatory interneuron (bGLNs) that plays an important role in the inhibition of the proboscis extension reflex that occurs in response to bitter gustatory stimuli.

## DISCUSSION

The experiments reported here identify a single pair of bilaterally homologous local interneurons in the SEZ of the adult fly. Anatomical, behavioral and functional data indicate that these interneurons receive specific synaptic input from bitter-sensitive GRNs. Moreover these data show that activation of the interneurons has a marked inhibitory effect on appetitive PER. From this, we conclude that the identified interneurons are first-order gustatory interneurons that have an important role in the aversive bitter-sensitive gustatory circuitry of the adult fly.

Previous studies have provided evidence for the notion that bitter and sweet gustatory stimuli are processed by different sensory interneuron populations in the SEZ of the adult fly, implying a segregation of the circuitry for aversive and appetitive taste modalities (Gordon and Scott, 2009). Several SEZ sensory interneurons involved in the circuitry for appetitive sweet taste processing have been characterized; these comprise both local and projection interneurons (Kain and Dahanukar, 2015; Yapici *et al.*, 2016; Kim et al., 2017). In contrast, previous to this report, little was known about sensory interneurons in the SEZ circuitry for bitter taste processing. Thus, the identification of this bitter sensitive gustatory interneuron in the of the adult fly represents a significant step towards understanding how bitter taste modalities are processed by the gustatory circuitry in the brain.

Our functional experiments involving targeted activation of the identified bitter-sensitive gustatory interneurons indicate that these cells are essential elements of the circuitry for aversive gustatory stimulus processing. Indeed, the strong PER response elicited by tarsal stimulation with high sucrose concentrations is all but eliminated by concomitant activation of these local interneurons. In wildtype animals, this type of powerful inhibition of PER responses is generally only seen if highly aversive and potentially toxic substances such as quinine or strychnine are mixed with the appetitive sugar stimulus (König *et al.*, 2014; French *et al.*, 2015). In this respect, the important and possibly vital role of the identified SEZ interneurons in bitter avoidance is underscored by experiments in which they are functionally inhibited. Remarkably in this case, the PER response to an otherwise aversive mixture of sucrose and quinine is significantly increased from the very low response level that occurs if the interneurons are functional to a level that could result in ingestion of toxic substance mixes in the wild.

VGN6341-based genetic access to the identified bGLNs sets the stage for future investigations focused on identifying their postsynaptic targets in the bitter gustatory circuitry of the SEZ. Several neuron types with neural processes in the SEZ that are known to be involved in gustatory/feeding circuitry might represent putative target neurons for these identified bGLNs. (Given that the first-order local interneurons have all of their neural arbors located within the SEZ, their target neurons must also arborize, at least in part, in the SEZ.) These include local interneurons with command function in the feeding program, projection interneurons that modulate feeding in the adult and are thought to combine feeding modulation with bitter taste detection in the larva, and motor neurons that control proboscis extension as well as food ingestion (Flood *et al.*, 2013; Hückesfeld *et al.*, 2015; Schwarz *et al.*, 2017). Whether these or other as yet unidentified SEZ neurons with roles in gustation or feeding are indeed postsynaptic targets of the fist order bitter-sensitive interneurons and whether they receive excitatory or inhibitory input from these cells must await further investigation.

It is remarkable that the bitter taste modality is conserved in insects and mammals, and that bitter gustatory information plays a key role in evoking aversive behavior in these and many other animals (Breslin and Spector, 2008; Yarmolinsky, Zuker and Ryba, 2009; Liman, Zhang and Montell, 2014; Joseph and Carlson, 2015). From this point of view, future studies of the structure, function and behavioral role of the bitter-sensitive gustatory circuitry in *Drosophila* are likely to be of general significance for understanding gustation in all animals including humans.

## MATERIAL AND METHODS

### Fly Stocks

*UAS*-*dTrpA1* (BL 26263 and BL 26264), *UAS*-*Shi^ts^* (BL 44222), *UAS*-mCD8::GFP (BL 5130 and BL 5137), UAS-CD8-RFP, LexAop-CD8-GFP (BL 32229), hsflp, UAS-mCD8-GFP (28832), Tub>GAL80> (BL 38880) were obtained from the *Drosophila* Bloomington Stock Center. VGN 6341GAL4 (Syed *et al.*, 2016). UAS GCamp6s, *G*r66a-LexA, lexA-op-CD4::SpGFP11, and UAS-CD4::SpGFP1-10 were provided by K. Scott (University of California); TshGAL80 by Julie Simpson (Janelia Farm Research Campus, VA, USA) (Clyne and Miesenböck, 2008)and Cha-Gal80 by T.Kitamoto (Kitamoto, 2002).

### Immunohistochemistry

Dissections of adult brains were carried out in 1X phosphate-buffered saline (PBS) and fixed in 4% freshly prepared PFA (prepared in 1 × PBS) for 30 minutes at room temperature. The fixative was removed, and the preparations were washed six times with 0.3% PTX (0.3% Triton X-100 in 1× PBS) for 15 minutes each at room temperature. Blocking of samples was performed for 15 minutes at room temperature in 0/1% PBTX (0.1% BSA in 0.3% PTX). Primary antibody diluted in 0.1% PBTX was added and samples were incubated at 4°C for 12hrs on a shaker. Primary antibodies used were chick anti-GFP (1:500; Abcam, Cambridge, UK), mouse anti-GFP (1:100; Sigma, cat# G6539), rabbit anti-HA (1:200, Abcam, Cambridge, UK) and mouse anti-nc82 (1:20, Developmental Studies Hybridoma Bank). Samples were washed in 0.3%PTX (15 minutes x 4 = 60 min). Fluorophore conjugated secondary antibody diluted in 0.1%PBTX was added and samples kept at room temperature for 2 hours on a shaker. Secondary antibodies (1:500; Invitrogen) conjugated with Alexa fluor-488, 568 and 647 were used in all immunostaining procedures. After removing the secondary antibody, samples were washed with 0.3%PTX (15 minutes x 4= 60min). To visualize muscle, rhodamine-conjugated phalloidin (1:200 Sigma) was used. Imaging was carried out using Olympus FV 1000 confocal point scanning. Z-stacks were collected with optical sections at 1μm intervals. Raw images were converted into TIFF files and exported to NIH Image J for image analysis. Further image processing was done using Adobe Photoshop.

### PER Assays

PER assays were done as described previously (Paranjpe *et al.*, 2012). Briefly, 0 to 12 h flies were transferred to a fresh food bottle for 1 d. Flies aged 2-4 days, 24-h starved, were used throughout the study. Each animal was mounted individually on a glass slide 4 h before assaying and kept in a humidified chamber to prevent desiccation. Immobilization of animals was carried out dorsal side down so that PER could be observed clearly under a stereomicroscope. Stimulation consisted of a tastant solution applied using a 1-mL syringe needle to the tarsal hairs. Animals were always water-satiated prior to tastant stimulation except in case of stimulation by water, in which flies were kept thirsty for 4h. Each stimulus lasted 2 sec. A test comprised five consecutive tarsal stimuli. A 10-sec gap was given between successive stimuli. Weak or completely failed PER events in response to a stimulus were scored as a zero. A successful PER response to a tarsal stimulus was scored as one. Thus, for example, a successful PER of 4 of 5 stimuli in a test would result in a performance score of 0.8.

### Clonal Analysis

GFP-labeled clones were generated in flies using transgenes to flip-out GAL80. Flies of the genotypes hs-flp, UAS-mCD8-GFP; UAS-dTRPA1, tub>Gal80>; VGN 6341-GAL4 were recovered from single heat shock treatment of 5-10 minutes at 37°C, which were performed at late third instar larval stage. Flies aged 3–6 days were tested in PER and immunohistochemistry assays as described.

### Calcium Imaging

Flies, 4-6 days old, were used for calcium imaging. Calcium imaging was done as described earlier (Harris *et al.*, 2015). Briefly, flies were anesthetized using CO2 and placed into a small slit on a custom-built plastic mount at the cervix so that the head was isolated from the rest of the body. The head was then immobilized using nail polish, and the proboscis was waxed at the maxillary palps in an extended position to allow for taste solution delivery. A coverglass was placed at the base of the rostrum at a 45° angle to the plane of the plastic mount so that the proboscis was isolated from the head. The head cuticle was dissected in ice-cold adult hemolymph-like solution (AHL) (Wang *et al.*, 2003) lacking calcium and magnesium, the air sacs were removed, and the esophagus was severed to allow viewing of the SEZ. Calcium- and magnesium-free AHL was replaced by AHL prior to imaging. Sweet stimulations were performed using 2 M sucrose, while bitter stimulations were performed with a solution containing 10 mM denatonium, 100 mM caffeine, 100 mM ATP, and 20% polyethylene glycol (PEG) (Harris *et al.*, 2015). To apply taste solutions to the proboscis, a glass pipette (with an outer diameter of 1.0 mm and an inner diameter of 0.78 mm) was filled with approximately 2 μl of taste solution using suction generated by a 1-ml syringe. The solution was drawn up the capillary slightly so that the tip of the capillary was empty. The capillary was then placed over the proboscis using a micromanipulator prior to image acquisition. At time point 4 of 20, slight pressure was applied to the syringe to deliver taste solution to the proboscis. Stimulations lasted approximately 4 s or the equivalent of three to four time points. There were 10-min intervals between successive stimulations. For heat activation of flies with genotype Gr66a LexA>LexAopTrpA1; VGN 6341> UAS-GCamp6s, IR laser was used to activate the Gr66^+^ bitter sensory neurons. All experiments were performed using an Intelligent Imaging Innovations (3i) spinning disk confocal system equipped with a 20× water objective. ImageJ was used to calculate minimum and maximum fluorescence intensities in selected regions of interest for each stimulus application, which was used to calculate ΔF/F.

## Author contributions

A.A.B. designed and performed most experiments, analyzed data, and co-wrote the manuscript. B.R.K. helped in Calcium imaging experiments. H.R. and K.V.R designed experiments, analyzed data and co-wrote the manuscript.

## Acknowledgements

We thank the *Drosophila* community and NCBS fly facility for generous supply of fly strains and antibodies. We are grateful to Kristin Scott UC, Berkeley and members of the Scott lab for helping with calcium imaging. We thank Sanjay Sane and Vatsala Thirumalai for helpful comments and discussions. This work was supported by Tata Institute of Fundamental Research and National Centre for Biological Sciences. We thank the Centre for Nanotechnology, NCBS (Department of Science and Technology Grant No. SR/S5/NM-36/2005), for the Olympus FV1000 microscopes in the Central Imaging and Flow facilities (NCBS). KVR was supported by J.C.Bose fellowship of the Department of Science and Technology.

## Supplemental Movies

### Movie S1, related to figure 1

The movie shows the response of VGN6341>TrpA1 flies before and after heat activation of neurons following stimulation on the leg with 10% sucrose

### Movie S2, related to figure 6A

**Bitter activates VGN6341 neurons in the Brain.** The movie shows sucrose-evoked changes of GCaMP3 fluorescence in VGN6341 neuron in the fly brain. Bitter was applied to proboscis at approximately frame 4. The dorsal is up.

### Movie S3, related to figure 6B

**Transgenic activation of bitter sensitive Gr66a^+^ neuron activates VGN6341 neurons in the Brain.** The movie shows changes of GCaMP3 fluorescence in VGN6341 neuron in the fly brain by transgenic activation of bitter sensory neurons by TrpA1. Bitter GSNs was activated at approximately frame 11. The dorsal is up.

